# Mice require proprioception to establish long-term visuospatial memory

**DOI:** 10.1101/2023.10.03.560558

**Authors:** Olivia Rutler, Serena Persaud, Stylianos Kosmidis, Jung M. Park, Nina Harano, Randy M. Bruno, Michael E. Goldberg

**Author notes:** Corresponding author, to whom questions should be addressed: Michael E. Goldberg. whose contributions are equal (Olivia Rutler, Serena Persaud). whose contributions are equal (Randy M. Bruno, Michael E. Goldberg).

## Abstract

Because the retina moves constantly, the retinotopic representation of the visual world is spatially inaccurate and the brain must transform this spatially inaccurate retinal signal to a spatially accurate signal usable for perception and action. One of the salient discoveries of modern neuroscience is the role of the hippocampus in establishing gaze-independent, long-term visuospatial memories. The rat hippocampus has neurons which report the animal’s position in space regardless of its angle of gaze. Rats with hippocampal lesions are unable to find the location of an escape platform hidden in a pool of opaque fluid, the Morris Water Maze (MWM) based on the visual aspects of their surrounding environment. Here we show that the representation of proprioception in the dysgranular zone of primary somatosensory cortex is equivalently necessary for mice to learn the location of the hidden platform, presumably because without it they cannot create a long-term gaze-independent visuospatial representation of their environment from the retinal signal. They have no trouble finding the platform when it is marked by a flag, and they have no motor or vestibular deficits.

## Introduction

Visual information enters the via the retina, but because the retina moves constantly the location of an object’s image on the retina does not necessarily correlate with the location of that image in space. The brain must transform the spatially inaccurate retinal signal to a spatially accurate signal, which is independent of the direction of gaze and which can be used for accurate perception and action [29]. Psychophysical evidence shows that head-fixed humans use two mechanisms for establishing visuospatial memory: (1) a rapid system that finds a remembered target after one or two saccades, and could arise from corollary discharge, and (2) a slow system used after more than five intervening saccades, which could arise from a stable craniotopic representation of the visual world [22]. There is a representation of eye position in area 3a of monkey somatosensory cortex which comes from sensors in the contralateral eye [31]. When the oculomotor region of area 3a is inactivated, a monkey can make a saccade to a remembered target after two intervening saccades but not after five or nine saccades [33], showing that somatosensory cortex provides the eye position information for long-term visuospatial memory. Here we asked if this finding were limited to the laboratory curiosity of long-term memory for saccades, or if proprioception were necessary for long-term visuospatial memory in general.

One of the salient discoveries of modern neuroscience is the role of the hippocampus in establishing gaze independent, long-term visuo-spatial memories. The rat hippocampus has neurons which report the animal’s position in space regardless of its angle of gaze [20], and rats with hippocampal lesions are unable to find a platform hidden in a pool of opaque fluid based on the visual aspects of their surrounding environment, the Morris water maze (MWM) [15]. We studied the effects of surgical lesions in the dysgranular region of mouse somatosensory sensory cortex (S1DZ), the equivalent of monkey somatosensory area 3a [26], on the ability of mice to learn the position of the hidden platform on the MWM, which specifically tests long-term visuospatial memory. We found that the mice were unable to learn the position of the hidden platform, although they did not forget the position of a partially learned platform. When we moved the platform to a different position, the lesioned mice were not able to learn the new platform location. The lesioned mice had no difficulty finding a new platform location marked by a flag, suggesting that proprioception is not needed for making the transition from a retinal to a spatial signal when the mice do not have to remember the spatial location of the goal. The mice did not have any motor deficit as assessed by their consistent swimming speed before and after lesions, or on the Rotarod test. They had no vestibular deficit as measured by time to turn in the air-righting test [18]. These results show that long-term visuospatial memory requires not only the hippocampal memory mechanism, but also the conversion of the retinotopic visual signal to a gaze-independent signal, a process dependent upon the eye, head and body position signals in somatosensory cortex.

## Methods

We first used 22 C57Bl/6 male mice and randomized them into lesion and sham groups.

The maze was a circular pool (120 cm in diameter) filled with water heated to 21°C and rendered opaque by 1 liter of nontoxic white tempura paint. The pool was surrounded by fireproof black curtains. In one quadrant of the pool there was a 14 cm wide hidden escape platform, upon which the mice could stand with their heads out of the water, but could not see. For the first two sessions we marked the hidden platform with a flag to accustom the mice to the maze. We then ran each mouse in the MWM four times each experimental session, placing each mouse in a different, pseudorandomly chosen starting quadrant of the maze. The mice learned the marked platform location in two sessions. We then removed the flag and mounted 4 large visual cues on the inner curtain wall of the maze where they remained unchanged throughout the duration of the experiment. The mice had to find the platform, which they could not see, by trial and error and use the visual cues to help them remember the platform’s location. We then ran the unoperated mice for 8 sessions. When we removed the flag, the mice did not remember the location of the platform. We recorded the mice’s swimming trajectory using an Edmund Optics video camera running at 250 frames per second, and extracted body position metrics using DeepLabCut. We measured the time from insertion into the maze until the time the mouse found the platform and stopped swimming. If the mouse failed to find the platform after 60 seconds, we guided the mouse to the platform with a finger, and allowed the mouse to sit upon the platform for 10 seconds before removing it from the pool. We used MATLAB and Excel for data analysis.

We performed the aspiration under anesthesia. Mice were maintained under isoflurane anesthesia at 2-3% and given carprofen 20mg/kg for analgesia. Craniotomies (2 mm x 0.5 mm) were made over the target dysgranular zones using stereotaxic coordinates (Paxinos & Franklin, 2019). Underlying cortical tissue was aspirated with a sterile blunt-tipped syringe connected to a vacuum (Figure 1A). Most lesions extended beyond the dysgranular zone boundaries into the barrel, shoulder, forelimb, hindlimb, and trunk regions of the primary somatosensory cortex (Figure 1B, C). An average lesion for each hemisphere was 0.53±0.03 mm^2^. In sham animals, craniotomies were made without removing the underlying tissue. Animals were allowed to recover for 3 days after surgery prior to behavioral testing. Lesions were objectively scored post hoc by experienced anatomists who were blind to the behavioral data. Lesions that extended into the hippocampus were excluded from analysis.

**Figure 1.**
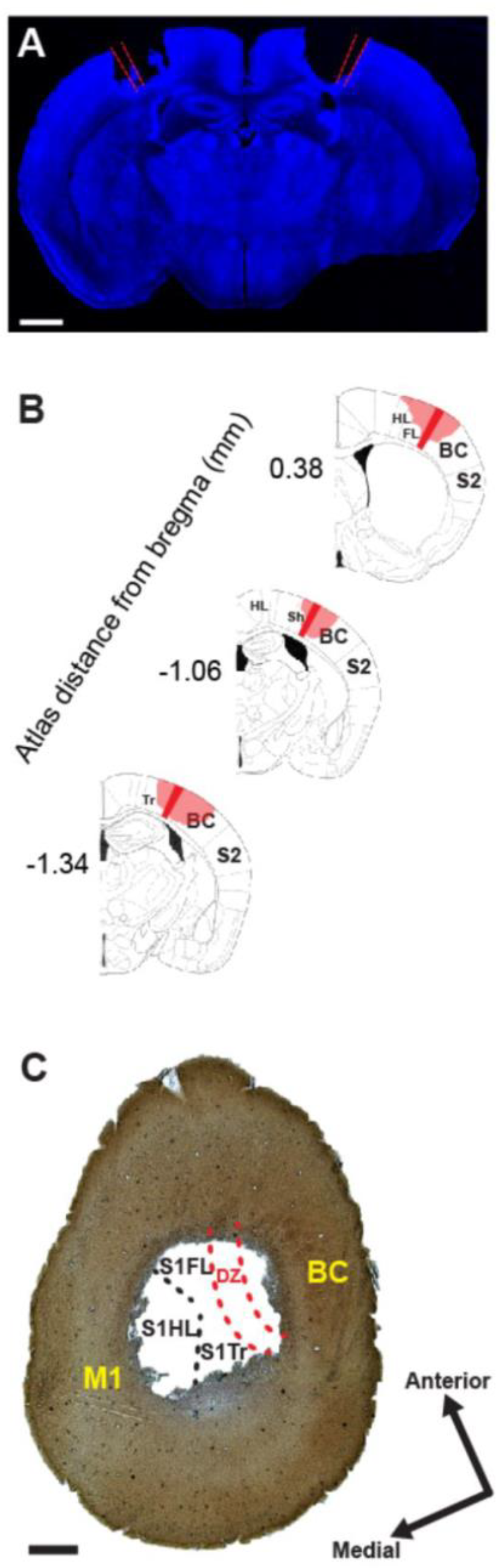
Bilateral dysgranular zone lesions. (A) Representative image showing extent of bilateral lesions.DZ boundaries in red. Blue,NeuN; scale bar 1mm.(B) Size and locations of DZ lesion in a mouse. The three locations along the anterior-posterior axis are shown overlaid onto a reference alias image from Pax inos & Franklin’s The Mouse Brain in Stereotaxic Coordinates, 5th edition.Numbers along anterior-posterior axis indicate approximate location rela tive to bregma.DZ boundaries in red,/e sions overlaid in pink.(C) Tangential section of cytochrome oxidase stained lesion.DZ boundaries in red. Yellow arrows indicating individual barrels of the barrel cortex. Scale bar 1mm. Abbrevia tions: BC -Barrel Cortex. DZ -Dysgran ular Zone. HI Hindlimb S1. FL forelimb SH Shoulder S1. TR Trunk S1. S1 Prima ry somatosensory cortex M1 Motor Cortex

Three days after the lesions, we ran the mice in the maze for another 8 sessions. The experimenters running the animals and analyzing the data were blind to whether a given mouse was in the sham or the lesion group. After the sham mice learned to reach the platform with the same latency that it took them to find the marked platform, we changed the location of the platform to a different quadrant of the maze, and measured the mice’s performance for another 5 sessions. Finally, to test whether the lesion affected the mice’s ability to find visual targets, we moved the platform to a different quadrant, marked it with the flag, and ran the mice for an additional session. To evaluate the mice’s vestibular system, we used an air-righting test: we dropped each mouse upside down and measured the time it took to right themselves. To evaluate their general motor ability, we measured the time they could stay on a Rotarod.

We perfused the mice intracardially with saline followed by 4% PFA in PB. We extracted the brains were extracted and incubated them overnight in 4% PFA. After washing three times in PB, we sectioned the brains coronally at 60µ or tangentially 100 µ thickness using a Leica VT1000S vibratome. We the sections permeabilized for 2 h with 0.4% Triton-X at room temperature and then incubated them overnight with primary antibodies at 4C diluted in 0.1% Triton-X in PBS. We washed the sections three times for 10 min in PBS prior to mounting them using flouromount mounting medium (Sigma-Aldrich). We acquired images using an inverted confocal microscope (Leica, LSM 700). For Nissl staining, we incubated coronal slices with Neurotrace 640/660 (1:200, ThermoFisher Scientific, cat#N21483). For cytochrome oxidase staining, we cut the cortex tangentially in 100 µm-thick sections. We quantified the lesion sizes were quantified

## Results

Inactivation of S1DZ impaired the mice’s ability to learn the location of the hidden platform using the environmental visual cues in the maze. When the platform was marked with a flag, all mice easily learned its location. Their latency decreased from 32 s on the first session to 11 s (p <0.0001, ttest) in the sham-lesioned group, and 28 to 12 s (p < 0.0001, ttest) in the S1DZ-lesioned group (Figure 2 left). When we removed the visual cue and moved the platform to a different quadrant, the mice gradually learned the location of the hidden platform (Figure 2, sessions 3-10). The future sham group’s time to escape fell from 34 to 20 s and the future lesion group’s time fell from 32 to 19 s. We then operated on the mice and tested them on the third session after the surgery to allow time for recovery and decrease chances of infection. The lesioned mice initially performed worse than they had just before the surgery, but they gradually improved such that, on the last three sessions, their escape times were not different from that of the last pre-lesion session (mean time on experimental sessions 10 and 20 were 18.9s and 17.2s respectively (p = 0.41, ttest). For the sham mice, the mean time on session 8, the session before the lesion, was 21.6 s; the mean time on session 20, the final session with the same platform location, was 11.4s (p < 0002, ttest). We then moved the platform to a new quadrant of the pool. The sham mice learned the new position quickly (difference between new platform session1 and session 5 (p < 0.0001) but the lesion mice did not (p = 0.28). Neither the lesion nor the sham mice had any difficulty finding a visually flagged new platform. On the flagged session the lesion group’s time was 11.0 s and the sham group’s time 11.8 s (p=0.72, ttest).

**Figure 2.**
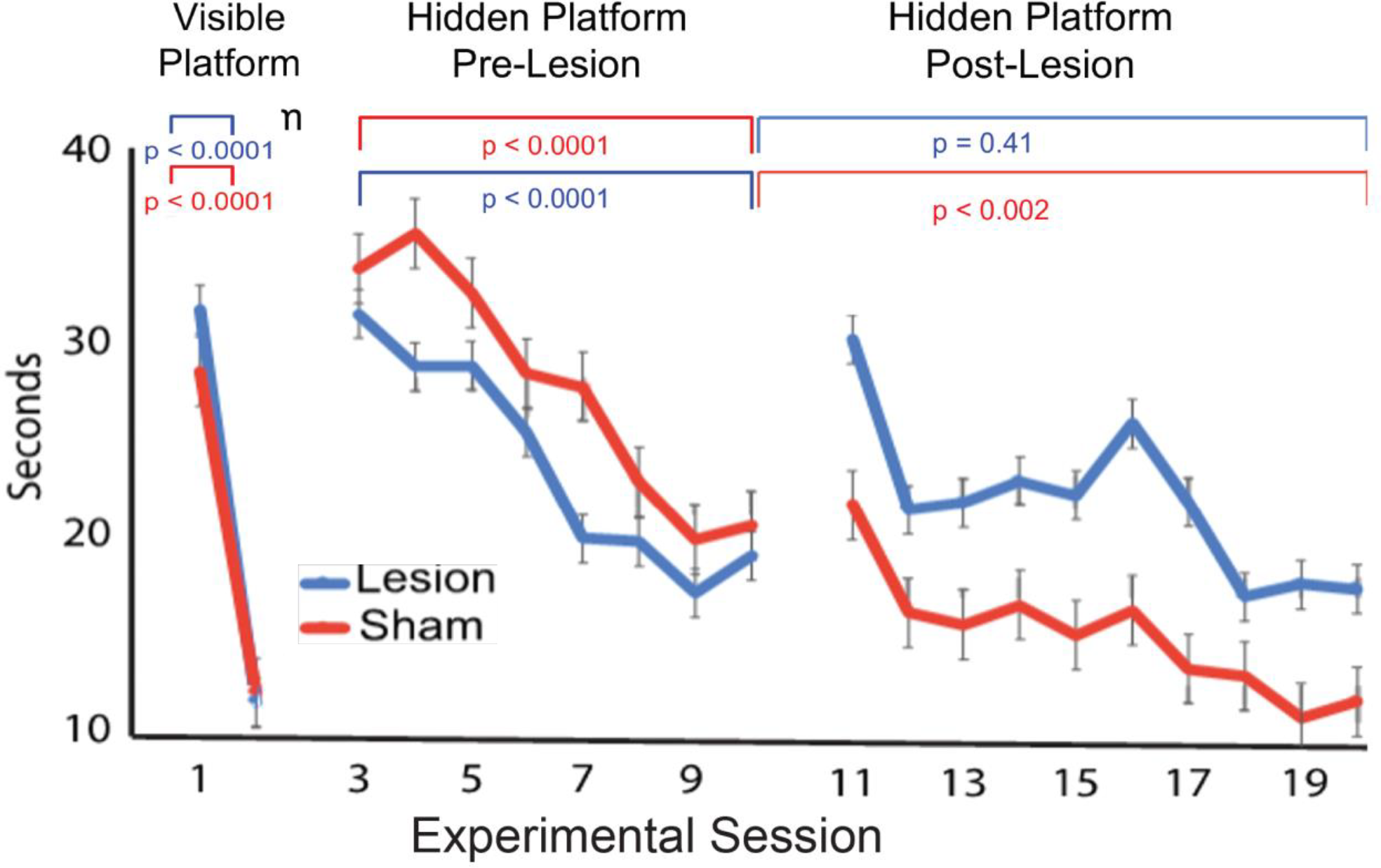
Mouse performance before and after lesion. Time to find platform (ordinate) otted against experimental day (abscissa). Left: flag visible. Center: Platform hidden, Platform hidden post pre-lesion. Right: Platform hidden post lesion. Error bars SEM.

**Figure 3.**
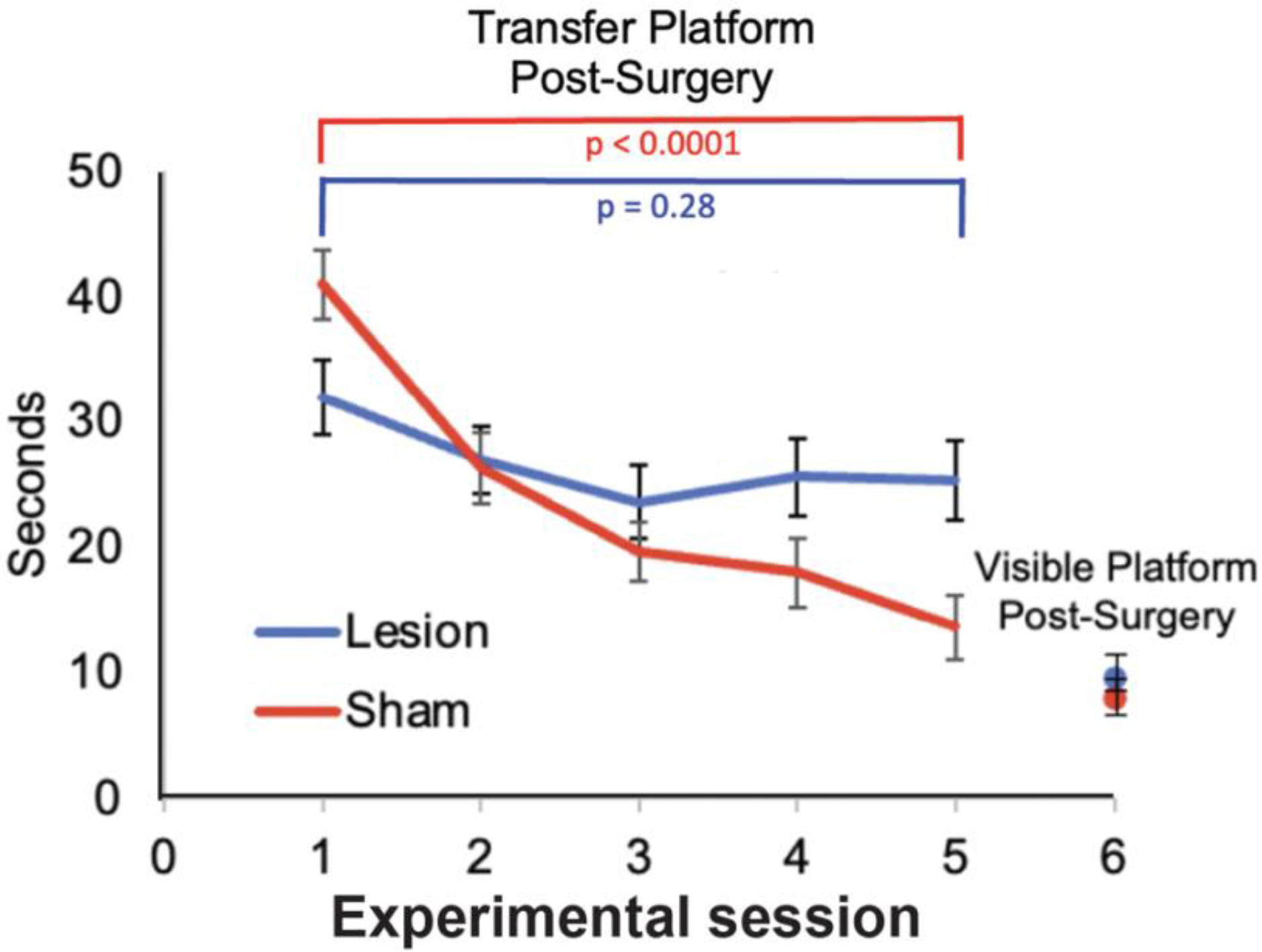
Lesion mice cannot learn a new platform position. Time to escape (ordinate) plotted against experimental day (abscissa) for lesion (red) and sham mice.

S1DZ in the mouse receives vestibular signals as well as proprioceptive signals [8]. Mice with genetically damaged vestibular systems exhibit a number of deficits, including an inability to swim [21], postural abnormalities [21] difficulty with air-righting and defective rotarod performance [18]. We tested air-righting in the two groups, dropping each mouse in a belly-up position onto a pillow 50 cm below for a total of four trials. The mean times from drop to righting were 394 ms for the sham mice, and 439 ms for the lesion mice, which were not statistically different (p = 0.31, t-test). We also tested rotarod performance: the lesion group trended towards better performance than the sham group: 10 out of 12 lesioned mice lasted for the maximum time of 300 s, while only 6 of 10 sham mice did so. Thus, our lesions did not produce gross motor deficits that could explain the performance deficits we observed.

## Discussion

In order to create a representation of visual space for perception and action, the brain must transform the spatially inaccurate visual signal transmitted by the constantly moving retina to a spatially accurate signal which is independent of the direction of gaze. The simplest example of this problem is the double-step saccade task (Supplementary Figure 1). In this task, a human or monkey has to make successive saccades to two stimuli, both of which appear and disappear before the first saccade. The retinal position of the first saccade determines the amplitude and direction of the first saccade. However, after the first saccade there is a dissonance between the retinal location of the target and the saccade necessary to acquire it, and the brain must compensate for this dissonance. Humans [9] and monkeys [7] perform the task easily.

Two solutions for this problem were proposed over 100 years ago. Hermann von Helmholtz suggested that the motor system solves the problem dynamically, feeding back the signal that will drive head and eye movements to the visual system to compensate for an impending movement [11]. Sir Charles Sherrington suggested that the brain measures the position of the eye in the orbit and the head on the shoulders, and uses these data to calculate the static position of an object in space [25].

Helmholtz saw a patient who sustained a paralysis of the lateral rectus muscle, the muscle that moves the eye towards the ear, in his only seeing eye. When the patient tried to look laterally the eye did not move, but the he perceived the visual world to jump in the opposite direction of the attempted movement and then slowly drift back to the resting position. This led Helmholtz to postulate that the motor system feeds back information about the impending saccade to upgrade the visual systems [11]. Today this feedback signal is called corollary discharge [28] or efference copy [12]. In support of Helmholtz’s theory, neurons in many areas, including the lateral intraparietal area (LIP), [5], the superior colliculus (SC) [30], the frontal eye field (FEF) [14], V3a [17], V4 [19], and the parietal reach area [3] remap their receptive fields around the time of an eye movement so that a stimulus, currently not in the receptive field of a visually responsive neuron, can excite the neuron when that stimulus will be brought into the neuron’s receptive field by an impending saccade. Monkeys with lesions of the medial dorsal nucleus (MD) of the thalamus cannot perform the double-step saccade accurately although they can make perfectly accurate visually guided saccades [13]. The MD receives a signal from the presaccadic burst neurons in the SC, and inactivation of the MD eliminates remapping in FEF neurons, suggesting that the SC provides the corollary discharge signal necessary for remapping, relayed to the cortex through the MD [27]. Humans with lesions in the parietal cortex cannot perform the double-step task when the first saccade is made into the visual field contralateral to the lesion, even when the target is in the ipsilateral, unaffected field [6,10].

In contrast, Sir Charles Sherrington discovered that the extraocular muscles have as many spindles, the muscle proprioceptive organ, as the lumbrical muscles in the hand. He suggested that the brain measures the position of the eye in the orbit and the head on the shoulders, and uses these data to create a spatially accurate representation [25]. In support of Sherrington’s theory, eye position modulates the responses of neurons in the parietal cortex to visual stimuli without changing the retinal location of their receptive fields, the gain field [1,2]. It is possible to calculate the position of the stimulus in craniotopic coordinates from gain fields [23,24]. Two possibilities for the source of the eye position signal have been proposed: corollary discharge of an intended eye position signal such as the one found on the extraocular motor neurons[2], and from direct measurement of the eye position. There is a representation of eye position in area 3a of monkey somatosensory cortex which comes from sensors in the contralateral eye [31]. However, the activity of Area 3a neurons lag the eye position by an average of 60 ms[32]. Furthermore, when a stimulus flashes in the receptive field of neuron in LIP for up to 150 ms after the monkey makes a conditioning saccade, the response is modulated by the presaccadic eye position, although the postsaccadic eye position provides the eye position signal when the stimulus flashes 250 ms after the saccade [32]. If gain fields were used to provide the spatial location of the second target in the double step task, one would expect that monkeys would be unable to perform the double step when the two stimuli are flashed 50 ms after a conditioning saccade, because the gain fields would provide an inaccurate spatial location of the target for the second saccade. In fact, the monkeys have no trouble performing the double-step task when the gain fields are inaccurate [32]. Finally, preliminary data suggest that monkeys require proprioception to make accurate memory-guided saccades after 5 intervening visually guided saccade (Zhang et al 2018).

Our results are consistent with the idea that mice, like humans and monkeys, utilize two mechanisms for transforming a retinal signal into a spatially accurate signal: 1) a rapid, corollary-discharge driven signal, which ensures that the representation of the visual world is accurate across eye and body movements even before the impending change of retinal position. If the mouse can see the flag, it can use corollary discharge to update the representation of the flag to compensate for its eye, head, and body movements and maintain a spatially accurate representation of the flag, the goal of its movements. The corollary discharge system would not be expected to be affected by a lesion in the somatosensory system. 2) a long-term, proprioception-driven signal, which establishes a gaze-independent representation of the visual world. This signal enables the mouse to use the visual world to develop a spatial representation of a location present in the world that the monkey can find but has never seen. We hypothesize that as in the monkey this representation is dependent upon the gain field mechanism, which requires proprioception (Zhang et al, 2018). The mice can find the visual platform because their short term, corollary discharge mechanism is still intact. We suggest that mice, like monkeys, have a gain field mechanism in the mouse equivalent of posterior parietal cortex, which uses proprioception to create eye-position modulation of visual receptive fields. They cannot remember the hidden platform’s location because their proprioceptive gain-field mechanism is damaged.

These results also suggest that the place cell system in the hippocampus is also dependent upon proprioception [20]. Without the hippocampus and its place cells the brain cannot construct a long-term memory of the visuospatial environment [15]. In order to create such a map, the hippocampus needs a stable, long-term, accurate representation of the visual world. It cannot rely on a retinal map, even one that compensates for eye and head movements. Our results show that proprioception is necessary to develop the map. It could be that the hippocampus itself uses proprioception to establish a gaze-independent visuospatial signal. It is more likely that the posterior parietal cortex, which projects to medial entorhinal cortex [4], provides the gaze-independent signal to the grid cells from which the hippocampus constructs the place cells [16]. Our finding suggests that place cells and grid cells themselves are affected by inactivation of S1DZ and that mouse parietal cortex has gain fields. Further work will be necessary to establish these hypotheses as facts.

## Acknowledgements

This work was supported by NIH grant 1 RO1 NS 113078-01, 1P30NEI 19007, and 1R01EY032938-01A1 (M.E. Goldberg, Contact PI), the Howard Hughes Medical Foundation (Eric Kandel), and NIH grants R01 NS094659 and R01 NS069679 (Randy Bruno). We are grateful for the electronic and computer support by the late Glen Duncan, machining support by Juan Caban, veterinary support provided by Dr. Rivka Shoulson and her team in the Department of Comparative Medicine, histological support by John Pellegrin, and administrative support by Whitney Thomas, Danielle Shank, Arthur Uhimov and Maria Gabriele.

---

**Supplementary Figure 1.**
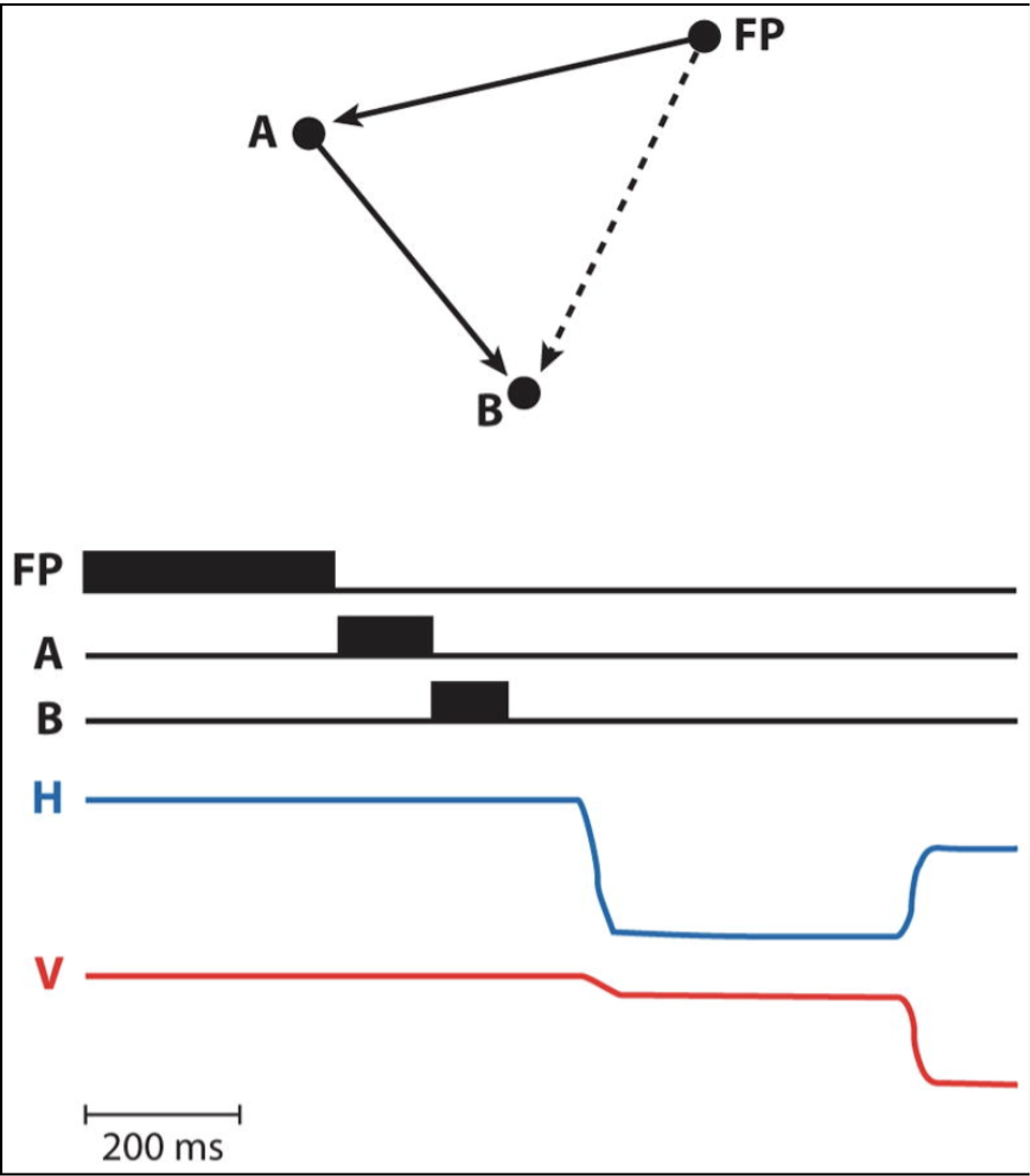
The double-step task [7]. A subject looks at a fixation point (FP), and two saccade targets, A and B, flash sequentially, each appearing and disappearing before the saccade begins (H and V, horizontal (left up) and vertical (left up) eye position. For the saccade to A the retinal vector FP-A is identical to the oculomotor vector FP->A. For the second saccade, the retinal vector FP->B is different from the saccade vector A->B, but can be calculated as the vector subtraction FP->B– FP->A.

